# Serotonin transporter knockout in rats modulates category learning

**DOI:** 10.1101/2020.11.09.373886

**Authors:** Chao Ciu-Gwok Guo, John Paul Minda, Judith Homberg

**Affiliations:** Donders Institute for Brain, Cognition, and Behaviour; Department of Cognitive Neuroscience, Radboud University Medical Center, Nijmegen, the Netherlands; Department of Psychology, The Brain and Mind Institute, Western University, London, ON, Canada

## Abstract

Lower function of the serotonin transporter (5-HTT) has a strong relationship with the development of autism spectrum disorder (ASD) in humans. One characteristic of ASD is the repetitive and restrictive behavior, which may form the basis for better memory and savant skills in some people with ASD. This characteristic in ASD may reflect a tendency towards an exploitation strategy rather than an exploration strategy during learning. Using a rat model, we developed a touchscreen-based task for testing 5-HTT knockout effects on stimulus category learning. By analyzing the data with a reinforcement learning drift diffusion model, we find that 5-HTT knockout rats show a lower learning rate and apply more of an exploitation versus exploration strategy compared to WT rats during category learning. The decision bound of decision-making during stimulus generalization indicates that more 5-HTT knockout rats than WT rats exploit irrelevant information to categorize stimuli. The touchscreen-based task we developed greatly increases the translational value from animals to humans and helps to understand the behavioral mechanisms underlying repetitive behavior in ASD.

## Introduction

Direct evidence from a human study showed that the functional efficiency of serotonin transporter (5-HTT) in the brain of people with autism spectrum disorder (ASD) is relatively lower compared to that of healthy control ^1^. Further, pharmacological inhibition of 5-HTT during the maternal period increases the risk for ASD in offspring ^2,3^, and inhibition of 5-HTT in children with ASD makes them even worse in repetitive behaviors ^4^. All these humane studies imply that lower function of 5-HTT has a strong relationship with the development of ASD.

A salient characteristic of ASD is repetitive and restrictive behavior ^5^. This behavior might be boring to most healthy people, but it seems interesting to people with ASD. It has been speculated that their repetition of things of interest is not strictly repetition for repetition’s sake ^6^. People with ASD are like domain experts, and are able to discriminate the subtle differences of objects they are interested in. The consequence of this ability is that one category of objects for most people, may be multiple categories for people with ASD. This allows people with ASD to have a novel experience in every “repetitive” behavior and be “restricted” to this behavior without getting bored. The “repetitive and restrictive” behavior may be the basis for many people with ASD that have better capacity of memory and form savant skills (also known as talent) ^7^.

Cognitive superiority and repetitive behavior have also been observed among individuals carrying the low activity short allele of the serotonin transporter-linked polymorphic region (5-HTTLPR), in both humans and monkeys ^8^. For example, short allele carriers show a better performance in reversal learning ^9^. Experiments in rodents also confirmed that the loss function of 5-HTT causes a better performance in reversal learning ^10,11^. One explanation is that ablation of 5-HTT increases cognitive flexibility. An alternative explanation might be that individuals characterized by reduced 5-HTT availability are better able to categorize the target stimulus and have a consistent approach to choose the target stimulus during reversal learning, namely through repetitive and restrictive behavior. Indeed, 5-HTT knockout (KO) rats show increased repetitive behavior when exploring objects and a developmental delay ^12^. The consistency of “repetitive and restrictive” behavior in ASD may reflect a tendency towards an exploitation strategy rather than an exploration strategy during learning ^13^. Exploitation refers to an explicitly chosen strategy to make decisions. In contrast, exploration refers to a strategy that might change or is not consistent when making a decision. During category learning, subjects enter a situation of exploitation-exploration trade-off.

5-HTT KO rats and people with ASD may have the capacity to learn categories correctly under rule-based instruction ^11,14,15^. However, people with ASD have a robust impairment in generalizing the learned information to novel situations/stimuli ^16,17^. Generalization is the process by which the brain processes information before taking actions. During this processing, the transfer of the stored information to novel situations/stimuli is required (Guo et al. 2020). Typically, the novel stimuli are similar to the categorized stimuli to some extent, such as the frequency of sound, tactile feel of the material, or orientation of gratings ^18^. According to the similarity relationship, one stimulus is associated with a valence (e.g. reward conditioned stimulus, CS), while the second stimulus is different from the first stimulus in one dimension (e.g. different orientation of gratings) and associated with another valence. The two CSs can be distinguished at the perceptual dimension. When using a series of stimuli (Generalization stimuli, GS) that vary along with the defined stimulus dimension (the orientation of gratings), generalization performance can be tested in rodents (Guo et. al. 2020).

In the current study, we investigated the effect of 5-HTT knockout in rats on exploitation and exploration learning strategies to categorize stimuli using a novel paradigm. In this paradigm, stimuli were presented on a touchscreen and subjects responded to stimuli directly by touching the screen. Presenting stimuli on the screen as targets is a widely used approach in human studies. Typically, disc-shape grating stimuli are used in humans to investigate rule-based category learning, in which subjects apply hypotheses to testing rules for forming stimulus categorization ^19–21^. We also investigated rule-based category learning in the present study employing rats as subjects. This increases the crucial translational value from animal to human studies. During the category learning task, the stimuli differed from each other on two dimensions, orientation and frequency. Subjects were required to focus on the orientation of the gratings while ignoring the frequency of the gratings in order to categorize the two preset categories correctly. In the orientation dimension, the standard deviation (SD) is the same in the two categories of grating stimuli, but the mean is different. The mean of stimuli is 25 in one category and 65 in the other category. In the frequency dimension the mean and SD parameters of the spatial frequency of the stimuli in the two categories are the same. People with ASD perform well in specific category learning but have a robust impairment in generalization ^16,17^. We therefore further tested whether 5-HTT knockout affects generalization in rats after sufficient training in stimulus categorization.

## Materials and methods

### Subjects

Eighteen male rats aged 95-110 days served as subjects. Nine of them were homozygous 5-HTT KO rats, the others were wild-type (WT) siblings. The sample size based on previous experiments ^11^(Guo et al. 2020). The KO rats (Slc6a41Hubr) had been generated by target-selected ethylnitrosourea induced mutagenesis and outcrossed for at least 15 generations with commercial Wistar (albino) rats ^22^. All rats were housed in a temperature-controlled room (21 ± 1 °C) with a humidity of 40-50% under a 12/12 h reverse light and dark cycle (lighting from 19:00 to 7:00). Rats were housed in pairs in regular European standard III H-type cages including a shelter. They had *ad libitum* access to water and food chow in home cages. Housing conditions and experiments were approved by the Animal Welfare Committee of Radboud University Medical Center, Nijmegen, the Netherlands.

### Stimuli

All stimuli used in the experiments were generated by using python (version 3.7) with the package of PsychoPy (version 3.0). Stimuli were presented on a touchscreen. The touchscreen is placed on one side of the wall of an operant box (more details about the operant box is described in Guo et al. 2020). The stimuli presented during category learning and generalization were black and white gratings of varying orientation (Orit) and spatial frequency (SF). Orit ranged from 0 to 1.5708 radians (rad) and SF ranged from 0.1202 to 0.4952 cycles per degree (cpd). These values are within the visual acuity of albino rats ^23^. Linear transformations normalized the dimensions of SF and Orit to create a two-dimensional space ranging from 0 to 90 cpd or rad. For example, 0.4952 cpd was converted to 90; 1.5708 rad was also converted to 90. The formulas for the conversion are as follows:

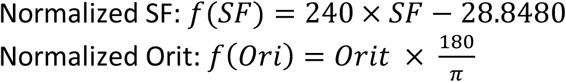

The diameter of the stimulus was set to 320 pixels so that the actual diameter distance of the stimulus was 6 cm presented on the screen. The specific stimuli parameters of SF and Orit dimensions during category learning and generalization are described below.

### Stimuli in category learning

The stimuli were divided into two categories during category learning. The boundary between the categories corresponded to the orientation of the alternating light and dark bands of the stimuli. One type of stimulus fell into Learning Category 25 (LC25) and the other into Learning Category 65 (LC65). All stimuli were sampled from a population of bivariate normal distribution, in which Orit and SF were the two variants. As shown in Figure 1a, points on the left side are members of LC25 and points to the right are members of LC65. The mean value of Orit is 25 degrees, and the SD is 2 for LC25 stimuli. The SD of Orit for LC65 is also 2, but the mean value is 65. In the dimension of SF, the mean values of SF for LC25 and LC65 stimuli are both 45, and the SDs are 18. The summary value of each category is as follows:

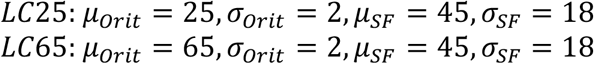

**Figure 1.**
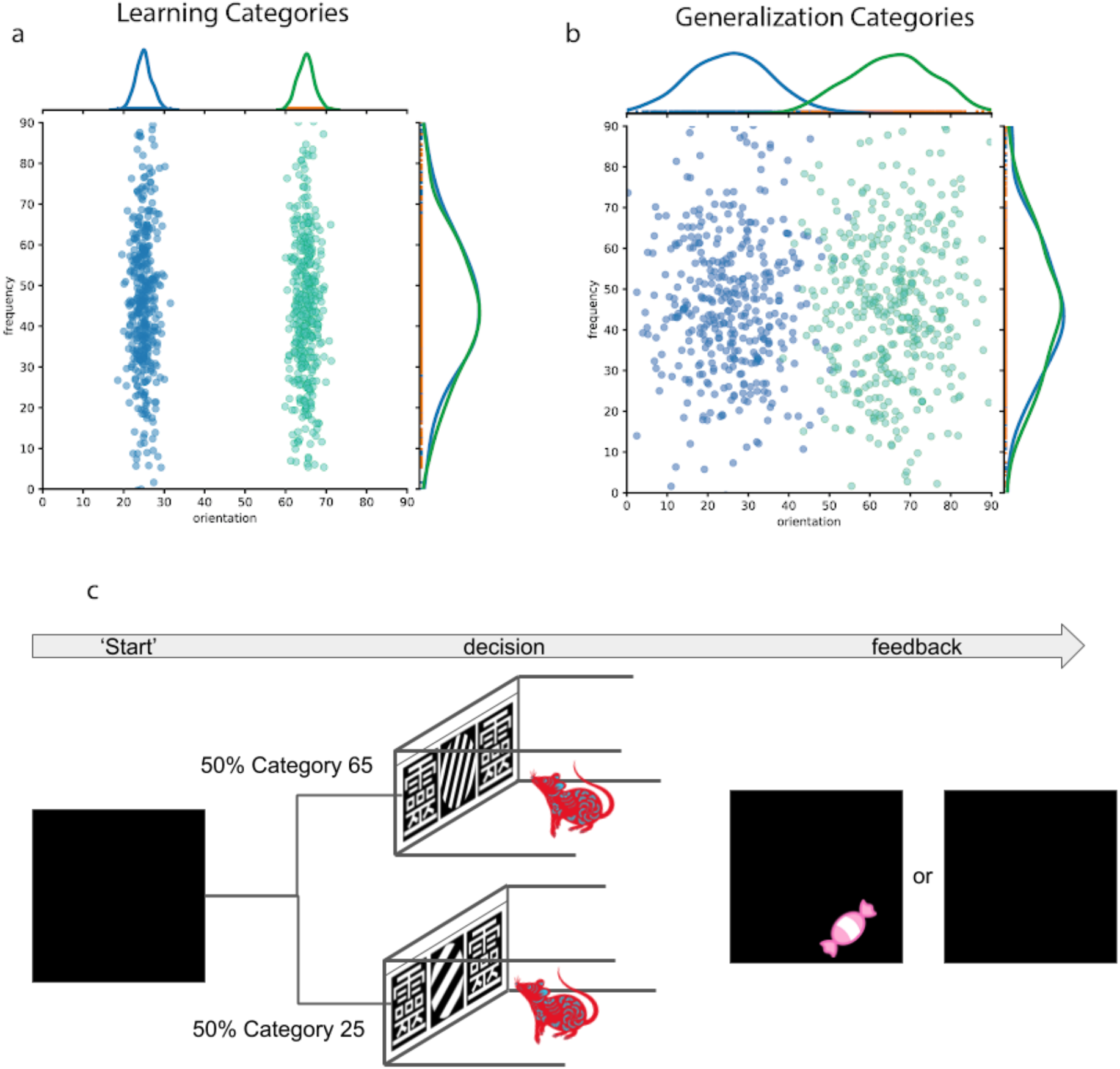
Distributions of stimuli and trial procedure. **(a)** The bivariate normal distribution of stimuli from the Learning Category 25 (blue) and learning Category 65 (orange). **(b)** The bivariate normal distribution of stimuli from the Generalization Category 25 (blue) and Generalization Category 65 (orange). **(c)** Trial procedure for category learning and generalization. If a subject assigns a stimulus to the corresponding category within a trial duration, a sucrose pellet is delivered as a reward.

### Stimuli in generalization

Stimuli presented during the generalization test had identical means of Orit and SDs of SF as the stimuli presented in category learning. The means along the Orit dimension were also identical, but the SDs were increased to 10. Figure 1b shows the stimuli distributions of both Generalization Category 25 (GC25) and Generalization Category 65 (GC65). The distribution parameters of the two stimuli sets were summarized as below:

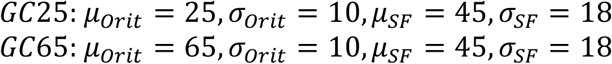

The test stimuli could be categorized into two stimulus types based on the Orit SD. The value of Orit within the two SDs of stimuli in Generalization Category are equal to the stimuli in category learning. These stimuli are termed ‘trained stimuli’. The remaining stimuli beyond the two SD ranges of Orit dimension are termed ‘generalization stimuli’.

### Experiment setup and task

#### Training stage

Rats were subjected to a simple instrumental learning task. The task included two stages: instrumental conditioning and sequential instrumental conditioning. Each learning stage had a passing criterion. Only when the rat reached the passing criterion it was allowed to proceed to the next learning stage.

##### Instrumental conditioning

Rats were allowed to freely explore the operant box. A white disc-shaped stimulus (size: 2.88 cm^2^) was displayed in the center of the touchscreen. When the stimulus was touched, rats received a sucrose pellet as reward immediately. Rats were trained during one session per day (except Sunday) until completing 70% trials during a session. Each session consisted of 30 trials, each trial was 30 seconds long, and the inter-trial interval (ITI) was 10 seconds. Touching the stimulus during each trial period was marked as completing a trial correctly. When 70% of the trial in a session was completed, the animal was allowed to enter the stage of sequential instrumental conditioning the next session.

##### Sequential instrumental conditioning

During this stage, target stimuli from the learning category were presented in the center of the touchscreen. If a stimulus was touched, a reporter stimulus (RS) was randomly presented on either the left or right side of the screen sequentially. The symmetric Chinese character “靈” served as RS. If the RS was touched, a sucrose pellet was immediately delivered. Rats were trained for one session per day (except Sunday) until reaching a criterion of 90% trials completed during a session. Each session consisted of 60 trials, 30 for stimuli in LC25 and 30 for stimuli in LC65. Each trial lasted for 30 seconds at maximum, and the ITI was random from 10 to 15 seconds. The order of displaying the LC25 and LC65 trials was random. The same trial would not appear more than 3 times consecutively. If the target stimulus and RS were touched during a trial period, the trial was marked as completed correct. If a rat completed at least 90% of the trials in a session, it would enter the stage of category learning the next day.

##### Category learning

Rats were subjected in this stage to one session per day except Sunday. Each session consisted of 60 trials, with 30 trials presenting LC25 target stimuli and 30 presenting LC65 target stimuli. Each trial lasted for 30 seconds at maximum, and the ITI was random, ranging from 10 to 15 seconds. The order of the trials presenting LC25 or LC65 stimuli was randomly assigned for each rat and for each session. The same type of trial did not appear more than 3 times consecutively. The difference from sequential conditioning was that when the target stimulus was touched, two identical reporter stimuli (RS, 靈) were presented on the left and right side of the screen at the same time. The representative trial sequence is shown in Figure 1c. If the target stimulus presented on the screen was from LC25, the rat would only receive a sucrose pellet as a reward after touching the RS of 靈 presented on the *left side* of the screen. No reward was obtained when the RS of 靈 on the *right side* was touched. In contrast, if the target stimulus was from LC65, the rat would only receive a sucrose pellet as a reward after touching the RS presented on the *right side* of the screen. Trials that ended with delivering a sucrose pellet were marked as completed correct. With this experimental design, rats were able to report to the experimenter a stimulus category in each trial. Rats were trained until reaching the learning criterion of completing at least 80% of the trials in a session correctly and across the last three consecutive sessions. Once the rat had reached criterion, it entered the next stage of stimulus generalization during the next session.

### Stimulus generalization

Rats were tested during one session per day. Each session consisted of 60 trials, of which 30 trials presented GC25 stimuli and 30 GC65 stimuli. Each trial lasted for 30 seconds at maximum, and the ITI was random ranging from 10 to 15 seconds. Each rat was tested for 5 sessions. The trial sequence was the same as the sequence applied during category learning except for that the LC25 target stimuli were replaced by GC25 stimuli and the LC65 stimuli were replaced by GC65 stimuli.

### Data analysis

#### Linear mixed-effect models

The total number of sessions needed for reaching criteria in the stages of instrumental learning, sequential instrumental learning and category learning were statistically analyzed using JASP (version 0.12) for t-tests. If normality of the data was violated, student t-test was replaced by the Mann-Whitey test. T-test could capture the general genotype effect on learning, to capture the session-by-session effect, linear mixed-effect regression modeling was run in software R (version 3.6.2) with package lme4 ^24^ for frequentist analysis, and brms ^25^ for Bayesian analysis. Genotype, session, and stimuli factors were entered as fixed effects. Subject was entered as a random effect, and the session was entered as a random slope, unless stated otherwise. Response time data were fitted in a log-normal distribution, while response choice data were fitted in a beta distribution. For frequentist analysis, the P-value was derived from Wald chi-square tests using the package *car* ^26^. P-values less than 0.05 were defined as significant. For Bayesian analysis, significant effects were calculated by 95% credible intervals (CrI) and the estimate (E) of the effect was given. If the 95% CrI did not include 0, the effect was deemed significant.

#### Reinforcement learning and drift-diffusion model analysis for discriminative conditioning

Trial-by-trial analysis could reflect the dynamic complexities of decision-making and learning during discriminative conditioning. To capture both the within- and across-trial dynamics, data were analyzed by a combined model of reinforcement learning (RL) and drift diffusion model (DDM), namely reinforcement learning drift diffusion model (RLDDM). The parameters in RLDDM were estimated in a mixed-effect Bayesian framework using the HDDM package (version 0.8) ^27^ with the HDDMrl module ^28^in python (version 3.6).

RLDDM included the drift rate scaling (*v*), decision threshold (*a*), non-decision time (t), positive learning rate (*pos_alpha*), and negative learning rate (*neg_alpha*) parameters. A higher drift rate scaling means an increased sensitivity to rewards or an increase in the degree of exploitation relative to exploration. A wider decision boundary results in a slower but more accurate decision, while a narrower boundary results in a faster but error-prone decision. A higher learning rate results in rapid adaptation to reward expectations, while a lower learning rate results in slow adaptation. The non-decision time parameter records the time spent on stimulus coding and movement processes ^29^.

Group-level parameters of the genotype effects were used to assess discrimination performance. The posterior density plots of each parameter are presented. We also report the posterior means (E) and 95% credible intervals (Crl). If the 95% CrI did not include 0, the effect was deemed significant. The proportion of the posteriors (P) in which the parameters for one genotype is greater/less than the other were examined. Model validation was assessed by posterior predictive checks, in which we checked by visual inspection whether the observed patterns of data were within the predicted range.

#### Decision bound analysis for generalization test

General recognition theory (or decision-bound theory) was used to estimate each rat’s decision bound during the stimulus generalization stage. In order to categorize stimuli, the General Recognition Theory (GRT) postulates that stimulus (perception) space is divided into multiple areas. These regions are separated by decision boundaries and category labels (such as the learned categories of LC25 and LC65). When observers regard the stimulus as falling within a specific area (LC25 or LC65), they act according to the learned categories. When the category is defined by a bivariate normal distribution (such as the distribution for LC25 and LC65), the optimal division of the perception area is a linear decision boundary ^30,31^. Within a trial, the distance between the stimulus and the decision bound determines the possibility of a response choice. Specifically, the decision bound is defined as:

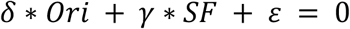

where Ori and SF are orientation and spatial frequency of a given stimulus, and δ, γ, ε are parameters. For the general linear classifier model, which is one of the models within the general recognition theory, the noise parameter (σ) is included. σ represents perceptual and criterial variance ^32^. Specifically, the subject's response choices were fitted in the model based with single orientation information (Ori model) and both spatial frequency and orientation (Ori-Fre model). Each model parameters were estimated using the maximum likelihood method and the goodness-of-fit statistic, Akaike information criterion (AIC),

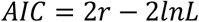

where *r* is the number of free parameters and *L* is the likelihood of the model given the data (^33^). The AIC statistic penalizes a model for extra free parameters in such a way that the smaller the AIC, the closer a model is to the true model regardless of the number of free parameters. If the best fit model is the one without spatial frequency information, it means that the subject’s response choice was controlled exclusively by orientation. The model was run in R with the package grt ^30^.

## Results

### 5-HTT KO and WT rats reached the criterion of (sequential) instrumental conditioning similarly

Rats touched the stimuli presented on the screen exploratorily during the instrumental conditioning. Independent t-test analysis revealed no significant difference between 5-HTT KO and 5-HTT WT rats regarding the numbers of sessions needed to reach the criterion of instrumental conditioning (t_(16)_=0.943, *p*=0.360, *d*=0.444, see Figure. 2a). An estimated Bayes factor (BF_01_ = 1.791) suggested that it was 1.791 times more likely that there was no genotype difference than there was genotype difference. During sequential instrumental conditioning, rats touched a target stimulus and subsequently a reporter stimulus to acquire a sucrose pellet. Mann-Whitey U test revealed that there was no significant genotype difference in reaching the criterion of sequential instrumental conditioning (U_(16)_ = 47.5, p = 0.467, r_B_= 0.173, see Figure. 2b). An estimated Bayes factor (BF_01_ = 1.835) suggested that it was 1.835 times more likely that there was no genotype difference than there was genotype difference during the sequential instrumental conditioning. Both BFs obtained in the stage of (sequential) instrumental conditioning are below 3, suggesting that the evidence for supporting no genotype differences is weak. Taken together, these analyses indicate that 5-HTT KO and WT rats likely associated touching the stimuli with acquiring rewards at a similar speed.

**Figure 2.**
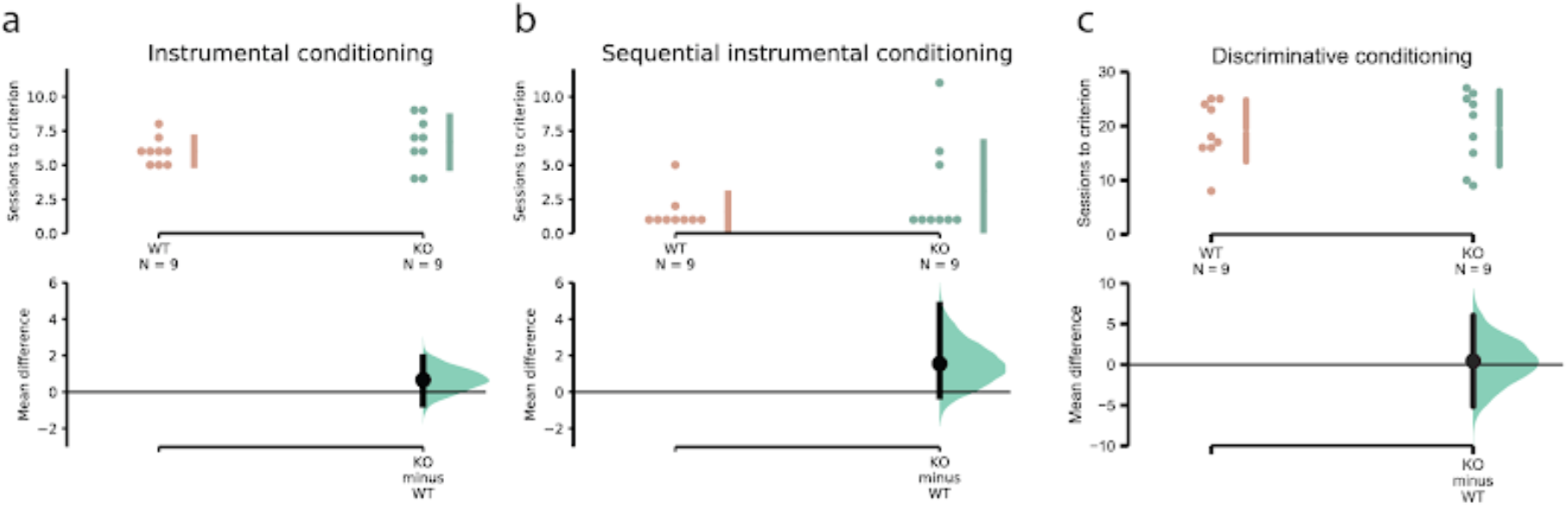
Number of sessions needed to reach criteria across different stages. Individual data are shown as dots. The effect size and 95% confidence intervals obtained from bootstrapping are plotted on separate axes beneath the individual data points. For each genotype, mean ± standard deviations are shown as vertical gaped lines. **(a)** Instrumental conditioning. **(b)** Sequential instrumental conditioning. **(c)** Category learning.

### Linear mixed-effect models reveal that 5-HTT KO and WT rats reached the criterion for category learning similarly

All rats were gradually able to categorize the two types of stimuli appropriately and reached the category learning criterion of at least 80% correctly across the last three consecutive sessions. An independent t-test showed that both KO and WT rats needed a similar number of sessions to reach the criterion (t_(16)_=0.150, *p*=0.883, *d*=0.071, see Figure 2c). An estimated Bayes factor (BF_01_ = 2.411) suggested that it was 2.411 times more likely that there was no genotype difference than there was a genotype difference for discriminative accuracy. Frequentist linear mixed-effect model analysis revealed that as the sessions progressed the percentage of correct response per session increased significantly (χ^2^_(1)_ = 158.7801, *p*<0.001). The effect of genotype and interaction between genotype and sessions were not significant (genotype: χ^2^_(1)_ = 1.6071, p = 0.2049; genotype*session: χ^2^_(1)_ = 0.8327, p = 0.3615). Bayesian linear mixed-effect analysis showed the same effects of session, genotype and genotype*session (genotype: E_(KO-WT)_ = −0.322, Crl = [−0.857, 0.288]; session: E = 0.11, Crl = [0.09, 0.14]; genotype*session: E = 0.01, Crl = [−0.04, 0.05], see Figure 3a). The mean response time (RT) of each session was also analyzed by a linear-mixed effect model. According to frequentist analysis the RT decreased significantly as the sessions progressed (χ^2^_(1)_ = 24.9077, p < 0.001). The effects of genotype and the interaction between genotype and session were not significant (genotype: χ^2^_(1)_ = 0.8535, p = 0.3556; genotype*session = 2.2893, p = 0.1303). When applying Bayesian analyses, similar results were found; the effect of session was significant, but no effect of genotype and interaction between genotype and session (genotype: E_(KO - WT)_ = 0.0961, Crl = [−0.26, 0.431]; sessions: E = −0.03, Crl = [−0.05, −0.02]; genotype*session: E = −0.02, Crl = [−0.05, 0.01], see Figure 3b) was found.

**Figure 3.**
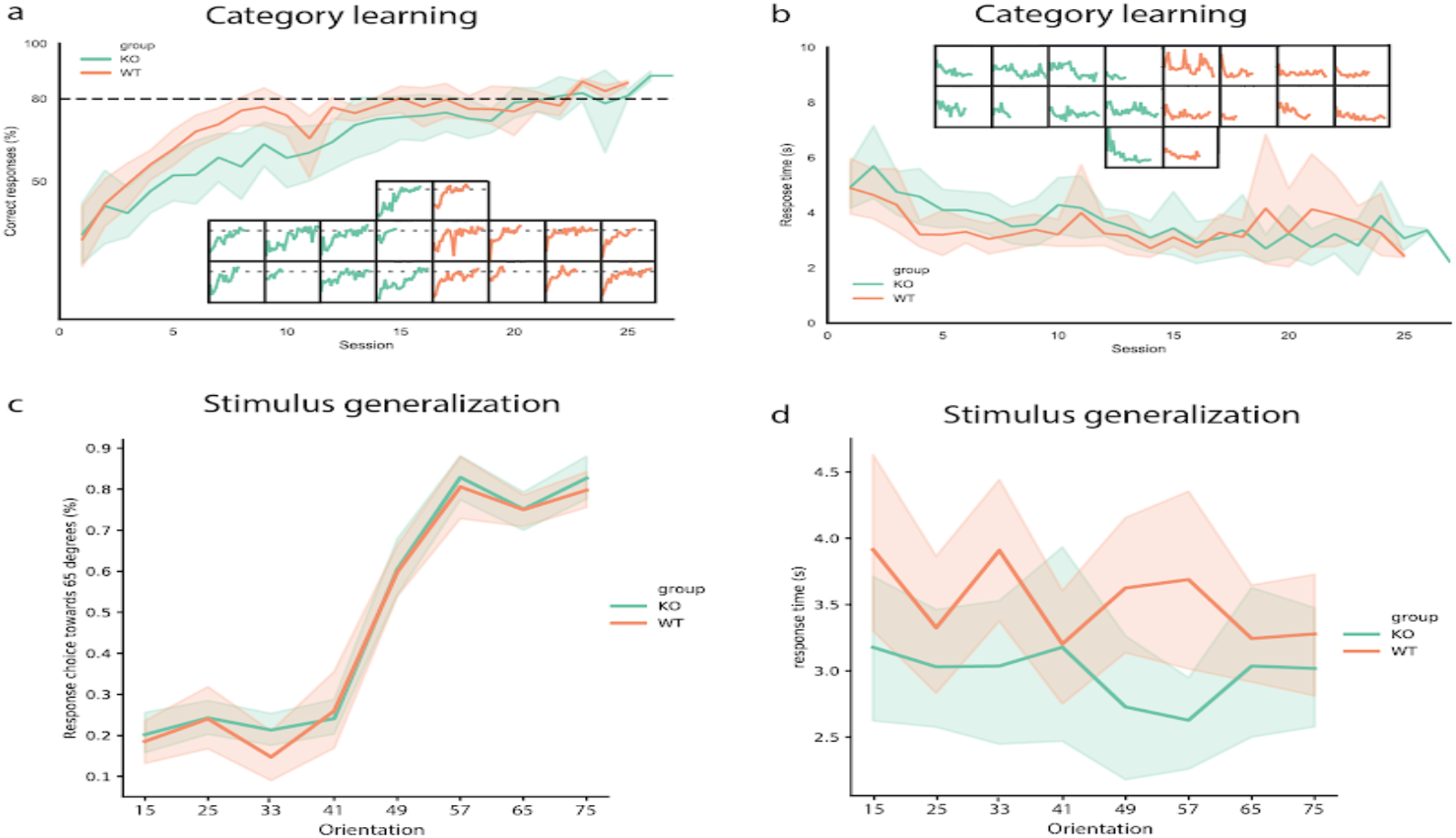
Linear Mixed-effect model analysis of category learning and stimulus generalization. Solid lines represent the mean of discriminative accuracy per session in each genotype. The transparent stripes are 95% confidence intervals. **(a)** The discriminative accuracy in each session. Figures embedded in panel **a** are the individual data of discrimination accuracy. **(b)** Response time (RT) in each session. Solid lines represent the mean of RT per session in each genotype. The transparent stripes are 95% confidence intervals. Figures embedded in panel **c** are the individual RT. **(c)** The percentage of response choice towards category 65 in each bin of orientation. **(d)** Response time in each bin of orientation.

Summarized, session-by-session analysis revealed that 5-HTT KO and WT rats reached the criterion in discriminative conditioning similarly.

### Linear mixed-effect models reveal that response choice and response time were similar between WT and KO rats during stimulus generalization

After reaching category learning criterion, rats were subjected to a stimulus generalization test. Frequentist linear mixed-effect model analysis revealed that 5-HTT KO and WT rats were not significantly different on the ratio of response choices (χ^2^_(1)_ = 0.0651, p =0.799). Bayesian linear mixed-effect analysis showed that there was no significant genotype effect for their response choice (E_(KO-WT)_ = 0.124, Crl = [−0.209, 0.484], see Figure. 3c). For response time, a frequentist linear mixed-effect model showed that KO rats responded slightly faster than WT rats (χ^2^_(1)_ =3.1556, p = 0.0757). However, there was no significant genotype effect on response time with Bayesian linear mixed-effect analysis (E_(KO-WT)_ = −0.215, Crl = [−0.505, 0.060], see Figure 3d)

### RLDD model reveals that KO rats had a higher drift rate scaling but lower positive learning rate than WT rats during category learning

To capture both the within- and across-trial dynamics, data were analyzed using the reinforcement learning drift diffusion model (RLDDM). The RLDDM was run with three chains with 40,000 samples (2000 was discarded as “burn-in”). Visually compared, the simulated data with the observed data from rats on both RT and choice proportion are decent (see Supplementary Figure 1a, 1b). A joint distribution of parameters is presented in Supplementary Figure 1c, showing that there was no obvious parameter collinearity in the model by visual inspection, supporting that the model was fitting the data well.

The RLDMM revealed that the posterior probability of drift rate scaling for the KO rats was significantly greater than that for the WT rats (see Figure 4 a1 and b1, E_(KO-WT)_ = 0.889, Crl = [0.123, 3.589]; d = 0.500, P_(KO>WT)_ = 0.993, P_(WT>KO)_ = 0.007). In addition, the posterior probability of positive learning rate for KO rats was less than that for the WT rats (see Figure 4 a4 and b4, E_(KO-WT)_ = −1.258, Crl = [−2.822, −0.392]; d= 0.226, P_(KO>WT)_ = 0.001, P_(WT>KO)_ = 0.999). No substantial evidence was found for genotype influences on the decision threshold (a), negative learning rate (neg_alpha), or non-decision time (t) (see Figure 4; *a*: E_(KO-WT)_ = 0.110, Crl = [−0.227, 0.445], d= 0.033, P_(KO>WT)_ = 0.755, P_(WT>KO)_ = 0.245; neg_alpha: E_(KO-WT)_ = 1.40, Crl = [−2.522, 5.004], d= 0.162, P_(KO>WT)_ = 0.792, P_(WT>KO)_ = 0.208; t: E_(KO-WT)_ = 0.013, Crl = [−0.160, 0.235], d = 0.040, P_(KO>WT)_ = 0.517, P_(WT>KO)_ = 0.483). The results above indicate that KO rats had a higher drift rate scaling but lower positive learning rate than WT rats.

**Figure 4.**
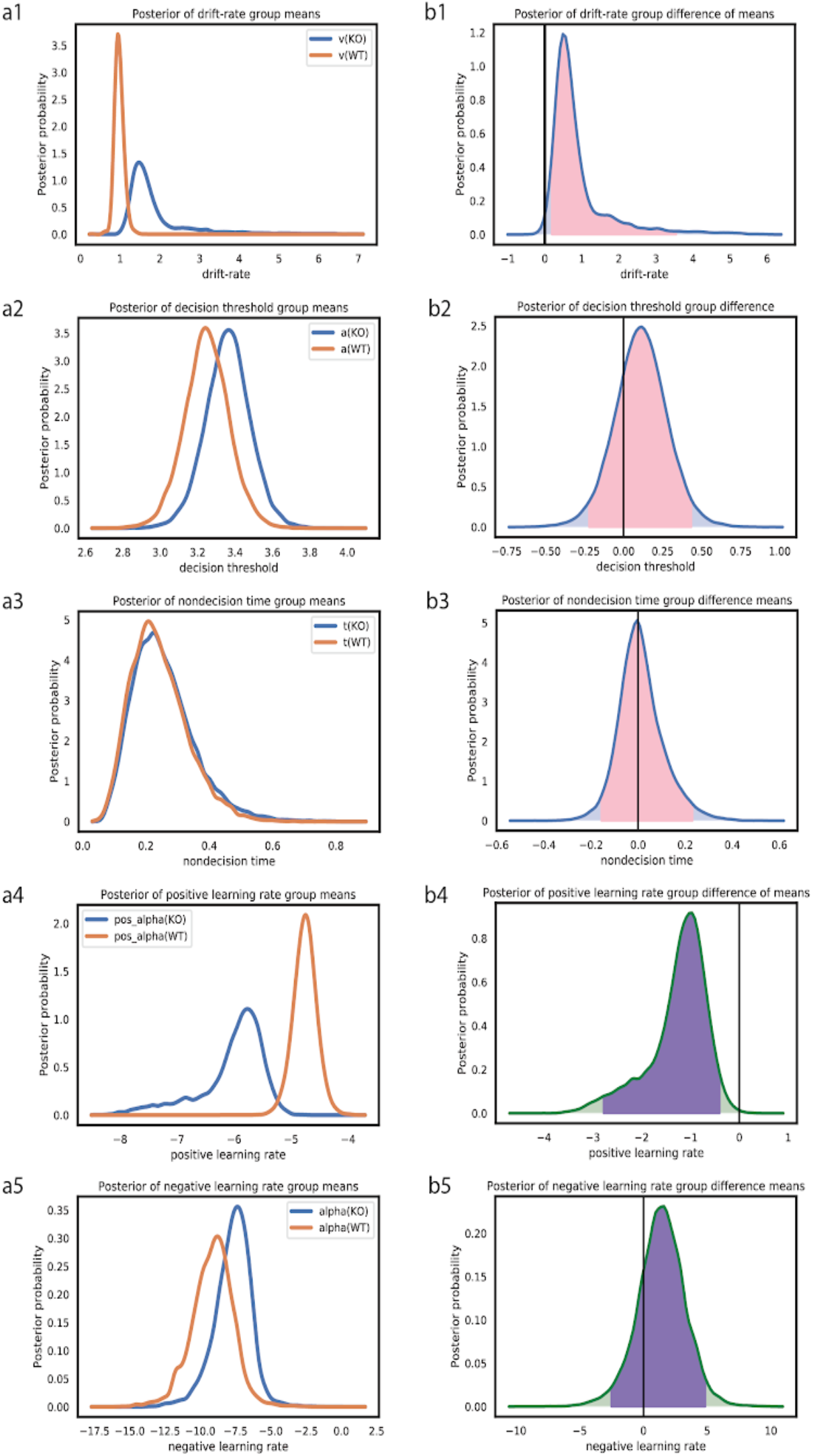
Reinforcement learning drift-diffusion model (RLDDM) analysis on category learning. **(a1-a5)** Density plots showing posterior distributions for WT and KO rats. **(a1)** drift rate scaling, **(a2)** boundary separation, **(a3)** non-decision time, and **(a4)** positive and **(a5)** negative learning rates. The learning rates were transformed by an inverse logit function to 0<alpha<1 for estimating normal distributions. **(b1-b5)** Posterior distributions of differences between KO and WT rats (KO-WT). Contrasts between KO and WT rats with at least 95% of the posterior distribution on either side of zero are considered significantly different. The 95% credible intervals are marked with shaded distributions. **(b1)** drift rate scaling, **(b2)** boundary separation, **(b3)** non-decision time, and **(b4)** positive and **(b5)** negative learning rates.

### More KO rats were interfered with irrelevant information during stimulus generalization

We used the general recognition theory to estimate the decision bound in generalization data (Figure 5a). Subjects’ responses were fitted with the model using orientation as information (Ori model) and also fitted with the model using both orientation and frequency as information (Ori-Fre model). Akaike’s information criterion (AIC) value was used to determine which model fitted to the subject's responses better. The proportion of subjects’ responses that best fit with the Ori model is shown in Figure 5b. 0.78 of KO and 0.89 of WT rats in session one; 0.67 of KO and 0.89 of WT in session two; 0.78 of KO and 1 of WT rats in session three; 0.89 of KO and 0.89 of WT rats in session four; 0.78 of KO and 1 of WT rats in session five were fit best by a model that assumed a single-orientation-dimensional rule. Mixed-effects logistic regression confirmed that there was a significant genotype effect on the type of best fit model (χ^2^_(1)_ = 3.848, p = 0.0498). The other effects were not significant (session: χ^2^_(1)_ = 0.5745, p = 0.448; session*genotype: χ^2^_(1)_ = 0.1328, p = 0.716). The results indicate that the responses of KO rats were less likely to be fit best by the Ori-model than WT rats. It could be that more rats in the KO group were interfered by the spatial frequency of stimuli, i.e., irrelevant information (spatial frequency), than WT rats.

**Figure 5.**
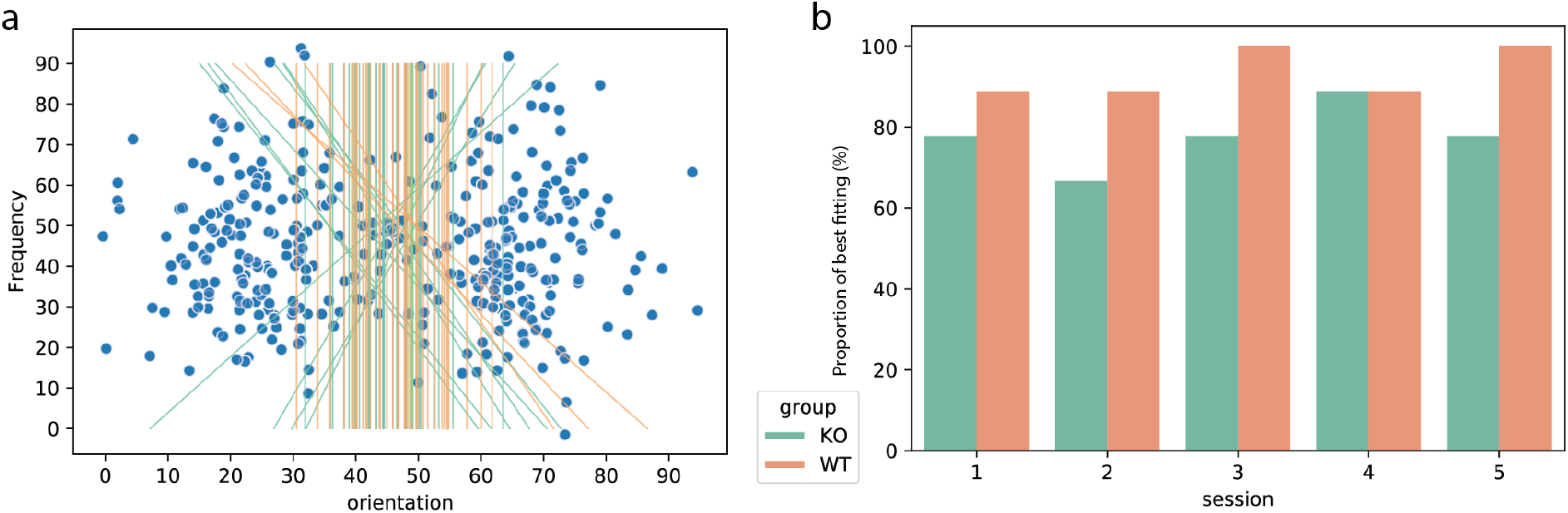
Decision bound analysis on stimulus generalization. (a) Estimated decision bound of each subject in the best fitted model. Blue dots represent orientation and frequency information of stimuli. The decision bounds of each rat of each session are denoted as the lines. (b) Proportion of subjects’ responses that best fit the Ori model in each session. A higher proportion indicates that more subjects favored making decisions based on orientation information and ignored the frequency information.

## Discussion

In the current study, we find that KO rats applied more of an exploitation versus exploration strategy to make decisions than WT rats. Furthermore, KO rats showed a lower learning rate than WT rats during category learning. This may be because KO rats focused more on stimulus details. Attention to detail could include attention to irrelevant features like the spatial frequency and might result in irrelevant information interfering with category decisions. This is consistent with our finding that during stimulus generalization more KO rats were worse in using only relevant stimulus information for generalizing choice-making.

During category learning, both KO and WT rats reached the learning criterion, indicating that KO 5-HTT in rats were able to categorize the stimuli successfully. This is consistent with the finding that people with ASD successfully learned to categorize stimuli (in a rule-based category task) similar to typical developed people ^15^. However, the prediction error and exploitation-exploration trade-off may affect category learning in people with ASD ^13,34^. For 5-HTT KO rats, a reinforcement learning drift diffusion model revealed that KO rats had a decreased learning rate for positive feedback compared to wild type rats. This indicates that KO rats adapted to reward expectations slower than WT rats. In addition, KO rats had a higher drift rate scaling than WT rats, indicating that the level of exploitation versus exploration in KO rats was higher than in WT rats. A reduction in 5-HTT availability may contribute to the development of ASD through the tendency towards a higher level of exploitation versus exploration trade-off during rule-based category learning. In addition, the repetitive behavior associated with reduced function of 5-HTT in people with ASD ^1^ may lead to a preference for an exploitation strategy. Further studies are needed to confirm such findings in humans.

People with ASD have been shown to exhibit an impairment in generalization ^16,17^. We found that KO rats tended to respond faster to stimuli than WT rats during the stimulus generalization test. Also, more KO rats than WT rats were likely interfered by irrelevant information during stimulus generalization. However, the majority of KO rats performed well in stimulus generalization. This indicates that 5-HTT KO rats may not have a robust impairment in generalization. The alternative explanation is that KO rats might form a memory trace where the reward irrelevant information (e.g. frequency of the stimuli) is linked to the reward relevant information (e.g. orientation of the stimuli). However, KO and WT rats’ accuracy of response choices were similar. In summary, 5-HTT KO rats might only have a mild impairment in stimulus generalization.

Serotonergic neurons are thought to signal the average reward information across trials ^35^. A recent study showed that serotonin neurons are activated by reward prediction errors during reversal learning ^36^. Further, learning the reward-predicting stimuli shapes the activity pattern of serotonin neurons gradually ^37^. All these findings indicate that serotonin plays an important role in reward related learning. In our current study, rats were learning to categorize stimuli to obtain a reward, in which 5-HTT KO rats (leading to elevated serotonin levels in the brain ^22,38^) reduced the learning rate compared to WT rats. During category learning, rats were facing an exploitation/exploration dilemma to correctly categorize the stimuli. To learn to distinguish the relevant (orientation) and irrelevant information (frequency), rats may explore all options to make decisions: a single source of information (orientation or frequency) or different sources of information combined (orientation and frequency joined). Rats may also exploit only one option to make decisions: orientation only, frequency only or orientation and frequency joined. The current study showed that 5-HTT KO rats applied more of an exploitation versus exploration decision strategy to categorize stimuli than WT rats did. This is in line with a recent finding that optogenetic activation of serotonergic neurons promotes an active exploitation strategy ^39^. Further generalization tests revealed that more KO rats' decisions were interfered by the irrelevant frequency information. The current model we use only reveals the decision bounds in rats during stimulus generalization. Other models are required to analyze generalization, such as whether KO rats have wide or narrow generalization ^18^. With narrow generalization, organisms might lack the ability to generalize the learned information to relevant situations; while with wide generalization, organisms might generalize the learned information to irrelevant situations.

The present findings may explain other phenotypes observed in the 5-HTT KO rats, such as reduced social interaction ^40^, lower ultrasonic communication during food exploration ^41^ and repetitive behavior ^12^, which are characteristic of ASD-like behaviors. More specifically, KO rats may have a reduced tendency to explore novel stimuli in their environment, and instead exploit stimuli they are familiar with, such as themselves (leading to less social interaction), previous food location and known objects, respectively.

Taken together, our study revealed that 5-HTT KO rats displayed more of an exploitation versus exploration strategy for decision during category learning. With this strategy, 5-HTT KO rats also displayed a mild impairment in stimulus generalization.

---

**Supplementary Figure 1.**
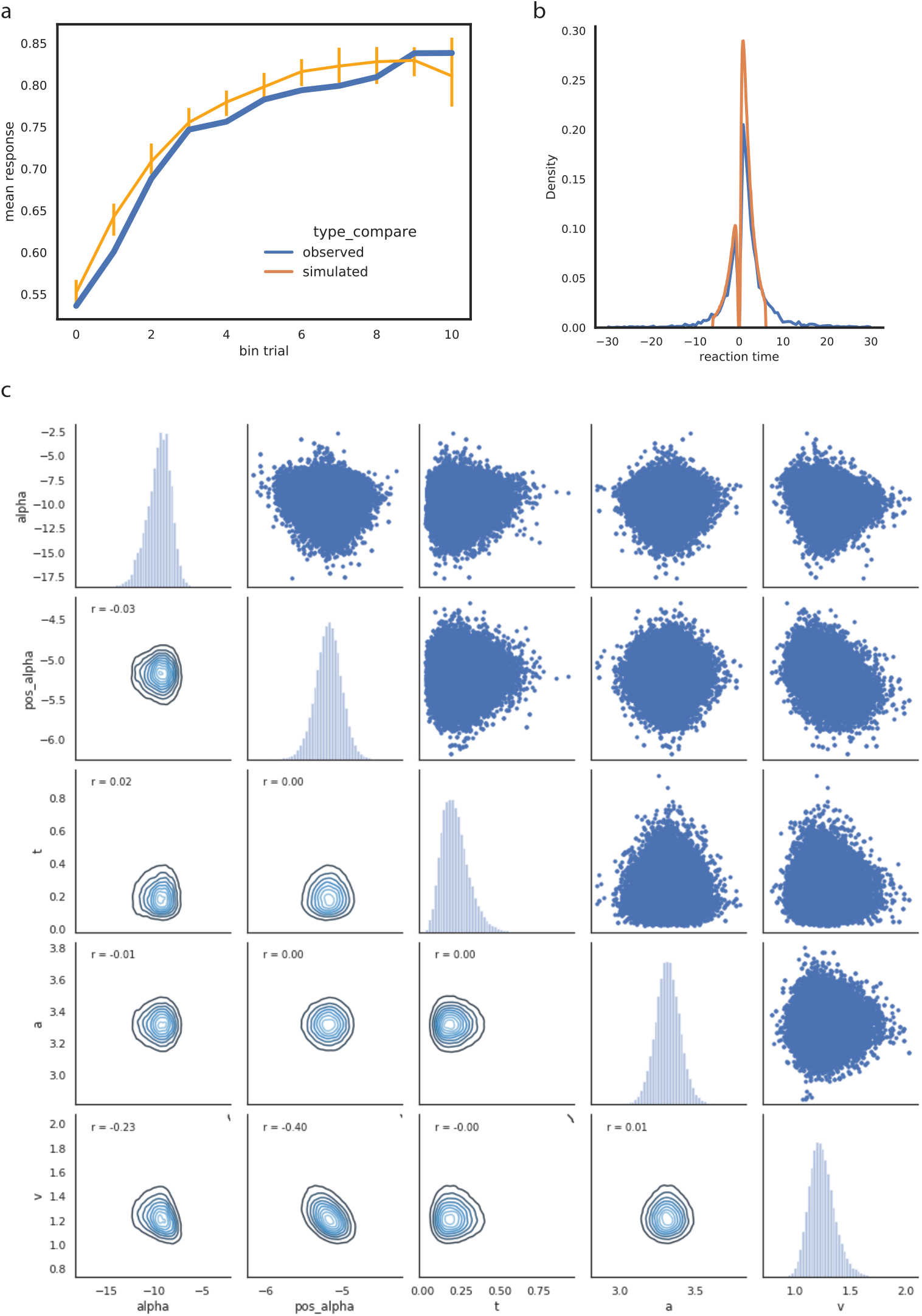
Posterior predictive check. **(a)** The plot shows the ratio of correctly classified stimuli throughout the learning process. The predicted data (yellow) closely follows the observed data (blue) except for over-predicting performance in the early phase. The uncertainty of the predicted data is captured by the 90% credible interval of the mean across the simulated datasets. **(b)** Density plot of observed and predicted response time (RT). The RT of the lower boundary selection (incorrect category selection) is set to negative so that the upper and lower boundary responses can be separated. **(c)** Scatter plot and density of group parameter estimates from posterior distributions. pos_alphs = learning rate for positive prediction errors (PEs), neg_alpha = learning rate for negative PEs, t = nondecision time, a = decision threshold, v = drift rate scaling.

## Acknowledgement

The work is financially supported by a scholarship of the China Scholarship Council awarded to CG (No. 201606140039).

